# Something to sink your teeth into: the mechanics of tooth indentation in frugivorous fishes

**DOI:** 10.1101/2024.10.05.616827

**Authors:** Jack Rosen, Karly Cohen, Cassandra M. Donatelli, Adam P. Summers, Stephanie Crofts, Matthew A. Kolmann

## Abstract

Frugivorous vertebrates engage in a mutualism with fruiting plants: the former receive a nutrient subsidy and the latter benefit by having their seeds distributed far from parent plants. Vertebrate frugivores like primates and bats have particular morphologies, like wide jaws and blunt teeth, which are thought to aid in dismantling fruit and obtaining trapped sugars. However, variation among frugivores and fruits has made the identification of common frugivore phenotypes difficult. We measured the performance of frugivorous fish dentitions whether this performance was comparable to fruit-eating bats and primates. We also explored how fruit characteristics affect puncture performance, and how indentation of fruit differs mechanically from harder foods like nuts. Finally, we used photoelasticity and videography to visualize how serrasalmid dentitions propagate stresses in simple gel models. We expected that frugivore dentitions would exhibit low force and then high work when engaging fruit tissues. Aligning with our expectation, the most frugivorous serrasalmid we tested, *Colossoma*, had dental performance that matched the low force, high work model. Indentation behavior differed between food types, both between fruits and nuts, and among different fruits. We also documented considerable differences in the indentation performances of different serrasalmid dentitions, among frugivores, omnivores, and carnivores. We propose that some differences in the morphology of frugivore dentitions make them better for granivory (eating seeds) than the softer fruit tissues. Fishes exhibit convergent mechanical and morphological strategies with other vertebrates for obtaining nutrition from fruits and seeds.

## INTRODUCTION

Fruit-eating vertebrates (frugivores) are a diverse lot – ranging from bats and birds to fishes and reptiles – and all participate in an ecological mutualism that structures plant communities (Fleming & Kress, 2016; Wotton et al., 2016). Flowering plants entice frugivores with appetizing, nutrient-laden fruits containing seeds that are later excreted (Milton, 1984; Jordano et al., 2011). By excreting seeds far from the parent plant, frugivores ensure that the plants establish new populations, reduce the chance that seedlings compete with their parent plant (Stiles & White 1986; Lambert, 1999), and limit the spread of disease in crowded communities (Augspurger, 1983). This mutualism is thought to be one of the catalysts for the diversity of flowering plants and their animal mutualists over the last 70 million years (Correa et al., 2007; Jordano et al., 2011; Rojas et al., 2012).

The phylogenetic, ecological, and morphological diversity of frugivorous vertebrates has made it challenging to find stable phenotypic commonalities, as much of this variation is likely driven by the specific mechanical challenges of fruits, seeds, and nuts (Kinzey & Norconk, 1990, 1993; Norconk et al., 2011). For example, some obligate frugivores may specialize on fruits from select plant species, or feed only on ripe fruits, while other frugivores are generalists, feeding on fruits as well as other plant or animal resources (Kinzey & Norconk, 1990; Dumont, 1999; Valenta et al., 2021). There are some generalizations about frugivores that can be made – in bats, for example – frugivores tend to have wide palates and skulls, and robust mandibles (Freeman, 1988, 1992; Dumont, 1997; Vogel et al., 2014). Frugivorous bats and primates also tend to have flattened dental surfaces adjacent to sharp cusps – an adaptation that may remove tough fruit rinds while keeping fruit pulp intact prior to processing (Freeman, 1988, 1992; Dumont, 1997). Research on the functional morphology of frugivores have focused on birds, bats, and primates; however, fishes are important frugivores in Neotropical systems and share some features in common with other fruit-eating vertebrates (Huie et al., 2019; Burns et al., 2021; Kolmann et al., 2024).

In the Neotropics, pacus (Figure 1), the herbivorous cousins of piranhas (Correa et al., 2007, 2014, 2015), play major roles as seed dispersers (ichthyochory). There are 60+ species of pacus, ranging from the largest (by weight) fish in the Amazon, *Colossoma macropomum*, a frugivore (Goulding & Carvalho, 1982; Roubach & Saint Paul, 1992; Lucas, 2008), to smaller-bodied taxa such as *Acnodon normani (*sheepshead pacu) that consume plant matter, insects, and even fish scales (Leite & Jégu, 1992). South American forest plants have co-evolved with pacus over the past 40-60 million years, alongside or even preempting other vertebrate seed dispersers (e.g., primates and birds; Correa et al., 2007; Kolmann et al., 2021). In fact, these fishes play such a crucial role in these flooded forests that overfishing poses a threat to overall rainforest functions (Correa *et al*., 2015a,b; Costa-Pereira & Galetti, 2015; Costa-Pereira et al., 2018; Araujo et al., 2021). Understanding phenotypic variation across vertebrate frugivores helps us understand whether and to what extent plant-frugivore mutualisms have shaped macroevolutionary outcomes.

**Figure 1.**
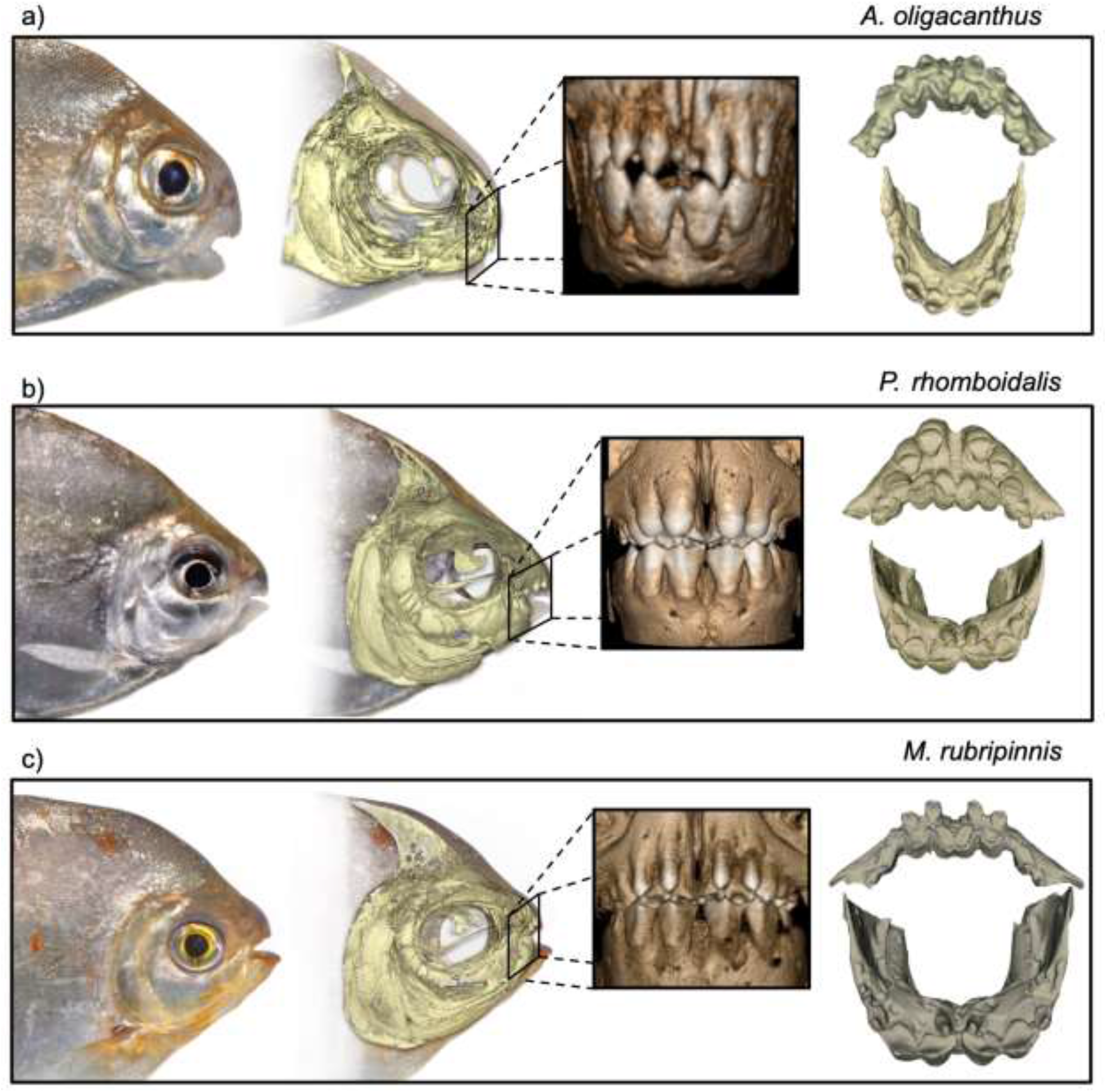
Cranial phenotypes of three pacu species used in this study. From left to right: lateral live images of three species of pacus (courtesy Oliver Lucanus), lateral ct scan volume renders, inset: anterior images of upper and lower dentition, upper (top) and lower (bottom) dentition of each species. (a) *Acnodon oligacanthus*, a basal myleine pacu and omnivore (Mol, 2012). (b) *Prosomyleus rhomboidalis*, a larger-bodied myleine pacu, omnivore and facultative frugivore (Boujard et al., 1990). (c) *Myloplus rubripinnis*, a generalized myleine pacu and omnivore (Dary et al., 2017).

What exactly do we expect from the performance of frugivorous fish phenotypes? Previous research modeling tooth function in frugivores has resulted in two heuristics we call the “High Force - High Work” and “Low Force - Low Work” models. The “High Force - High Work” model (Figure 2A), also known as the Mortar & Pestle, stems from studies on bats and primates (Lucas, 1979; Lucas & Luke, 1984; Kubitzki & Ziburski, 1994). This model states that frugivore teeth should effectively pulverize fruit tissues from the initial grasping of the fruit through to prey processing (i.e., when teeth are fully engaged in fruit tissues). In this scenario, the crushing and shearing actions of blunt teeth are used to burst as many plant cells as possible, releasing juice in the process (like a mortar and pestle). A lateral shearing motion, accomplished with the mobile mandibular joint of mammals, is a key to this model. The alternative relies on cutting teeth, and is best described with a “Low Force, Low Work” model (Figure 2B): tools with pointed tips minimize force of initial puncture and bladed edges minimize work during crack propagation (Abler, 1992; Crofts et al, 2020). We expect this scenario to characterize the performance of teeth in carnivorous piranhas like *Serrasalmus* (Berkovitz & Shellis, 1976; Kolmann et al., 2019).

**Figure 2.**
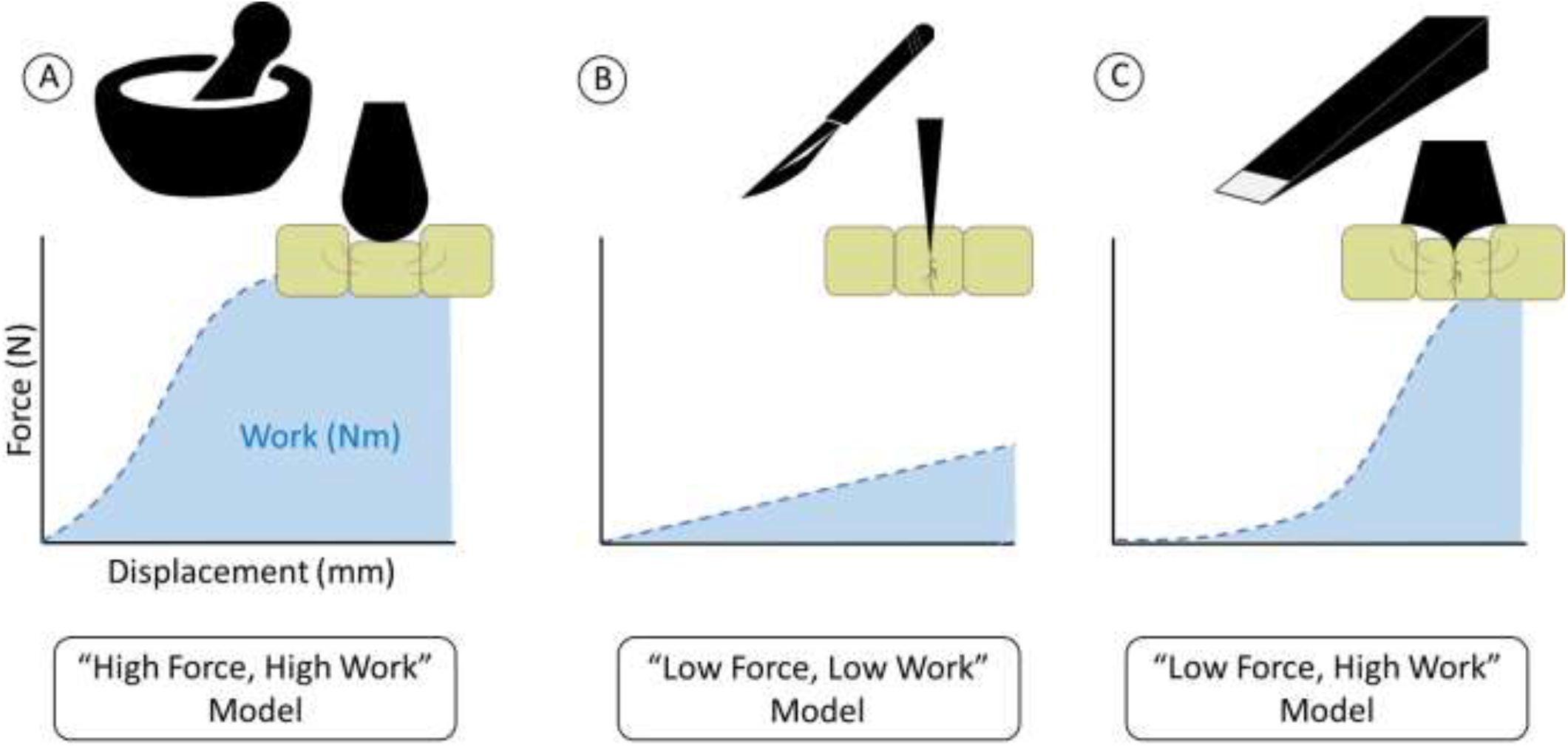
Hypothetical expectations for fish tooth performance. (A) Mortar & Pestle or “High Force, High Work” model. (B) Slicing Tool or “Low Force, Low Work” model, both characteristics of vertebrate frugivore dentitions. (C) Gripper & Pestle Tool or “Low Force, High Work” model in carnivores.

Alternatively, we propose that frugivorous fish tooth performance might be described best by what we call the “Gripper and Pestle” or “Low Force, High Work” model (Figure 2c). In this scenario, frugivore performance is facilitated by an initial piercing event (for gripping fruit) with a narrow, central tooth cusp, followed by a more crushing bite as broader, lateral tooth cusps maximize the contact area between the tooth and plant cell walls (Lucas & Luke, 1984; Freeman, 1988; Crofts et al., 2020). This mechanism allows for fine adjustments such as the ability to separate edible pulp from inedible rind (Freeman, 1988; Dumont, 1999), akin to a melon baller. Many pacus are bobbing for fruit from below and maintaining purchase on the fruit is of paramount importance.

Frugivorous fishes contribute to the dispersal of 25% of riparian plant species in some stretches of the Amazon rainforest; yet the feeding mechanics these fishes use to access seeds in fruit are entirely unexplored (Goulding, 1980). We used CT scans of pacu and piranha specimens to make 3D models of their dentitions, and then used these models to indent a variety of fruits and seeds. This study had four objectives: (1) Quantify the force (compressive loading) & work required to puncture and propagate cracks (indentation) in different fruits using a universal testing system, (2) compare puncture and indentation performance of different serrasalmid species’ dentitions, and (3) describe how stress is propagated by different dentitions using gel visualization. Finally, we (4) evaluate which of the three hypotheses for frugivore tooth function best describe pacu puncture performance. We expect that the tooth indentation performance of the most frugivorous pacu, *Colossoma*, will most closely resemble the “Low Force, High Work” model we proposed above (Figure 2). From the gel visualization of indentation tests, we expect that the blunt teeth of *Colossoma* will dissipate stresses over a broad area, rather than concentrating stresses at the cusps. This will contrast with carnivore dentitions, like in *Serrasalmus*, where points of high stress will be localized only at tooth crowns until the teeth are deeply embedded in the material.

## MATERIALS & METHODS

### Specimen selection and computed tomographic scanning

We selected six species of serrasalmids that span the gamut of diet variation, from piscivores and omnivores to obligate frugivores (see Nico & Taphorn, 1988; Nico, 1991; Röpke et al., 2014; Kolmann et al., 2021). *Colossoma macropomum* is a specialist frugivore (Goulding, 1980; Goulding et al., 1988), followed by *Prosomyleus rhomboidalis*, which consumes fruit in addition to a wide variety of other plant products (leaves, flowers, stems), as well as occasional insects (Boujard et al., 1990; Andrade et al., 2019). *Acnodon oligacanthus* consumes a wide variety of prey, but still is biased towards a plant-based diet (Planquette et al., 1996; Mol, 2012). *Serrasalmus rhombeus* are carnivorous opportunists, either actively hunting or scavenging for food (Nico & Taphorn, 1988). Finally, we included the taxon *Pristobrycon maculipinnis* - a poorly known but at least partially frugivorous piranha. Nico (1991) described the diet of this taxon (as “*Pristobrycon* sp.”) before it was formally described by Fink and Machado-Allison (1992). Its phylogenetic position is still unknown and we continue to refer to it as “*Pristobrycon*,” despite the genus’ current defunct status (Kolmann et al., 2020). We rendered 3D surface models from micro-computed tomography scanned museum specimens (see Table 1). Specimens were μCT scanned using either the Bruker Skyscan 1173 at Friday Harbor Labs (Bruker microCT, Kontich, Belgium) or the Nikon XTH 225ST (XTek, Tring, UK) at the University of Michigan’s Museum of Zoology. Scan parameters for each specimen can be found in Table 1. Raw CT image stacks were reconstructed, exported as .bmp or .tiff stacks, and then imported into 3DSlicer (slicer.org; Fedorov et al., 2012), for segmentation and surface rendering according to the workflow outlined by Buser et al. (2020).

**Table 1a.** Cherries, Load at puncture. Analysis of Variance (ANOVA) results for indentation tests using cherries. Significance was considered with an alpha greater than or equal to 0.05.

|  | diff | lwr | upr | adj. p-value |
| --- | --- | --- | --- | --- |
| *Myloplus-Colossoma* | 3.362 | 1.210 | 5.514 | 0.000 |
| *Prosomyleus-Colossoma* | 11.350 | 9.197 | 13.502 | 0.000 |
| *Acnodon-Colossoma* | 1.920 | -0.232 | 4.072 | 0.108 |
| *Pristobrycon-Colossoma* | 3.704 | 1.552 | 5.856 | 0.000 |
| *Serrasalmus-Colossoma* | -0.925 | -3.078 | 1.227 | 0.809 |
| *Prosomyleus-Myloplus* | 7.987 | 5.835 | 10.139 | 0.000 |
| *Acnodon-Myloplus* | -1.443 | -3.595 | 0.710 | 0.377 |
| *Pristobrycon-Myloplus* | 0.341 | -1.811 | 2.494 | 0.997 |
| *Serrasalmus-Myloplus* | -4.288 | -6.440 | -2.136 | 0.000 |
| *Acnodon-Prosomyleus* | -9.430 | -11.582 | -7.278 | 0.000 |
| *Pristobrycon-Prosomyleus* | -7.646 | -9.798 | -5.494 | 0.000 |
| *Serrasalmus-Prosomyleus* | -12.275 | -14.427 | -10.123 | 0.000 |
| *Pristobrycon-Acnodon* | 1.784 | -0.368 | 3.936 | 0.162 |
| *Serrasalmus-Acnodon* | -2.845 | -4.997 | -0.693 | 0.003 |
| *Serrasalmus-Pristobrycon* | -4.629 | -6.781 | -2.477 | 0.000 |

**Table 1b.** Cherries, Work to puncture. Analysis of Variance (ANOVA) results for indentation tests using cherries. Significance was considered with an alpha greater than or equal to 0.05.

|  | diff | lwr | upr | adj. p-value |
| --- | --- | --- | --- | --- |
| *Myloplus-Colossoma* | -1.183 | -6.231 | 3.864 | 0.983 |
| *Prosomyleus-Colossoma* | 1.475 | -3.572 | 6.522 | 0.956 |
| *Acnodon-Colossoma* | 2.485 | -2.562 | 7.532 | 0.705 |
| *Pristobrycon-Colossoma* | 23.369 | 18.321 | 28.416 | 0.000 |
| *Serrasalmus-Colossoma* | -0.956 | -6.003 | 4.091 | 0.994 |
| *Prosomyleus-Myloplus* | 2.659 | -2.389 | 7.706 | 0.642 |
| *Acnodon-Myloplus* | 3.668 | -1.379 | 8.716 | 0.287 |
| *Pristobrycon-Myloplus* | 24.552 | 19.505 | 29.599 | 0.000 |
| *Serrasalmus-Myloplus* | 0.228 | -4.820 | 5.275 | 0.999 |
| *Acnodon-Prosomyleus* | 1.010 | -4.038 | 6.057 | 0.992 |
| *Pristobrycon-Prosomyleus* | 21.893 | 16.846 | 26.941 | 0.000 |
| *Serrasalmus-Prosomyleus* | -2.431 | -7.478 | 2.616 | 0.724 |
| *Pristobrycon-Acnodon* | 20.884 | 15.836 | 25.931 | 0.000 |
| *Serrasalmus-Acnodon* | -3.441 | -8.488 | 1.606 | 0.358 |
| *Serrasalmus-Pristobrycon* | -24.324 | -29.372 | -19.277 | 0.000 |

**Table 1c.** Cherries, Work to max load. Analysis of Variance (ANOVA) results for indentation tests using cherries. Significance was considered with an alpha greater than or equal to 0.05.

|  | diff | lwr | upr | adj. p-value |
| --- | --- | --- | --- | --- |
| *Myloplus-Colossoma* | 0.548 | -15.652 | 16.748 | 0.999 |
| *Prosomyleus-Colossoma* | 2.420 | -13.780 | 18.620 | 0.998 |
| *Acnodon-Colossoma* | -12.598 | -28.799 | 3.602 | 0.219 |
| *Pristobrycon-Colossoma* | -19.013 | -35.213 | -2.813 | 0.012 |
| *Serrasalmus-Colossoma* | -43.929 | -60.129 | -27.729 | 0.000 |
| *Prosomyleus-Myloplus* | 1.872 | -14.328 | 18.072 | 0.999 |
| *Acnodon-Myloplus* | -13.146 | -29.346 | 3.054 | 0.180 |
| *Pristobrycon-Myloplus* | -19.560 | -35.760 | -3.360 | 0.009 |
| *Serrasalmus-Myloplus* | -44.477 | -60.677 | -28.277 | 0.000 |
| *Acnodon-Prosomyleus* | -15.018 | -31.218 | 1.182 | 0.085 |
| *Pristobrycon-Prosomyleus* | -21.432 | -37.632 | -5.232 | 0.003 |
| *Serrasalmus-Prosomyleus* | -46.349 | -62.549 | -30.149 | 0.000 |
| *Pristobrycon-Acnodon* | -6.414 | -22.614 | 9.786 | 0.857 |
| *Serrasalmus-Acnodon* | -31.331 | -47.531 | -15.130 | 0.000 |
| *Serrasalmus-Pristobrycon* | -24.916 | -41.116 | -8.716 | 0.000 |

### Surface rendering, model preparation, and 3D printing

We used a combination of segmentation tools to digitally dissect pacu and piranha dentitions (upper and lower jaws, as well as their teeth) from the rest of the cranium in 3DSlicer (Fedorov et al., 2012). We used 3DSlicer’s ‘wrap solidify’ tool with 14 iterations to make surface render models without any holes or surface imperfections, before exporting models as .stl files. We used MeshLab (Cignoni et al., 2008) to attach rectangular platforms to the upper and lower surfaces of the upper and lower jaws, respectively. These models were then exported to PreForm (3.26.0, Formlabs, Somerville, MA), where models were oriented to the same plane, scaled to a common size (4.0 cm wide), at the rearmost or most lingual tooth position of the lower jaw. We 3D printed our models using Grey v4 resin with 0.05 mm layer thickness in a FormLabs Form2 printer (FormLabs, Form2, Somerville, MA). Finished models were rinsed in a 99% isopropyl bath and then cured using UV light for 80 minutes (FormLabs FormCure, Somerville, MA).

### Mechanical testing

We chose to use whole dentitions, rather than isolated teeth, since piranhas and pacus regularly reorient prey and prey will engage with many teeth during each feeding. These behaviors typically only engage the anterior-most set of teeth (Cohen et al., 2023; Kolmann, unpublished data), so testing was done with fruit placed between these teeth. We used Gorilla Tough & Clear double-sided mounting tape to affix the models to the mechanical loading frame, and jaws were aligned to mimic their natural pattern of occlusion (Kolmann et al., 2019; Figure 3).

**Figure 3.**
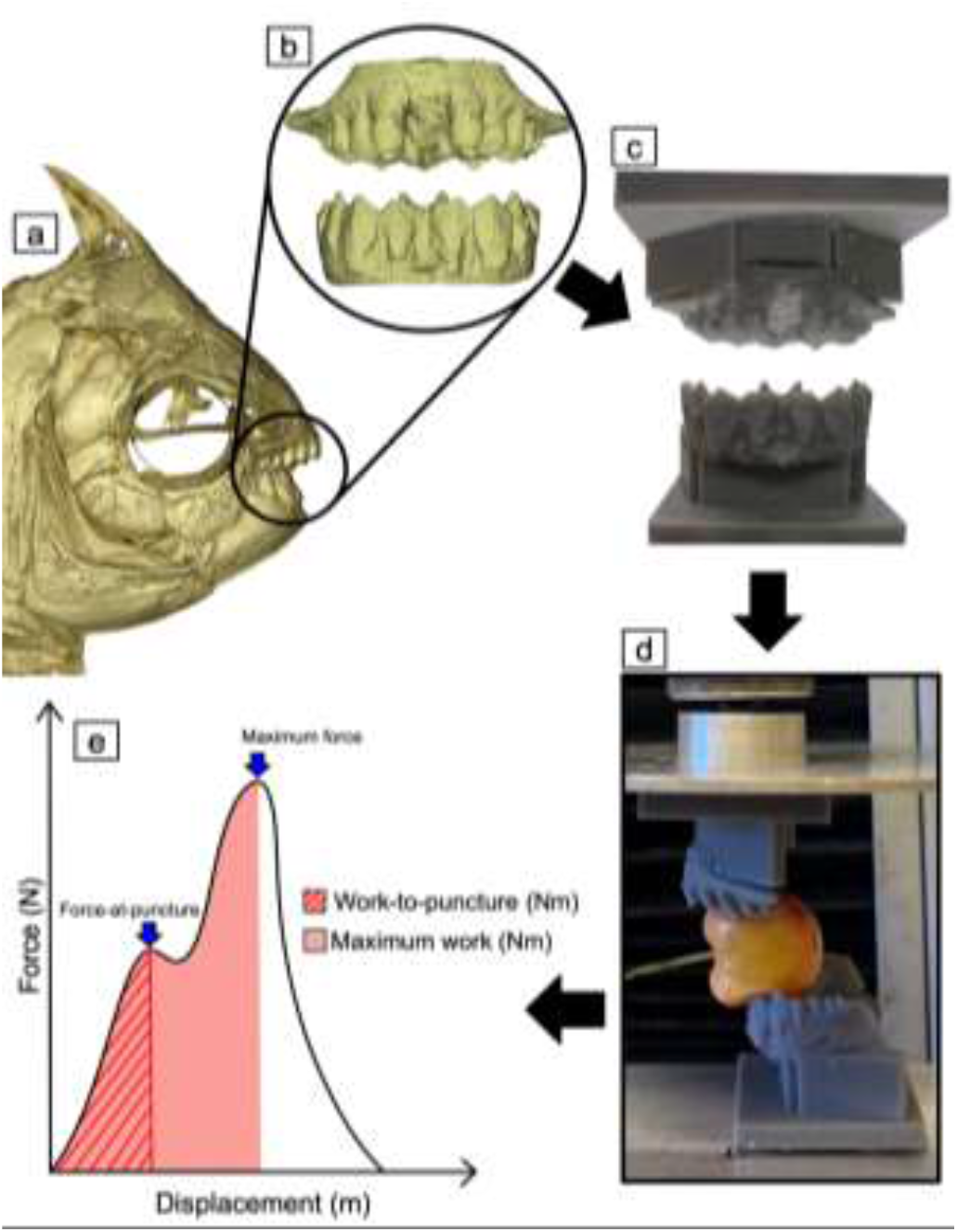
Experimental apparatus and representative force-displacement curve. (A) µCT scan of *C. macropomum* skull, (B) segmented dentition after wrap solidify and (C) 3D printed model of the dentition. (D) Photograph of mechanical loading frame with 3D resin-printed teeth affixed. (E) Example force-displacement diagram with relevant data points (Load-at-puncture, Work-to-puncture, Maximum Force, Maximum Work) highlighted. Work was calculated as the area under the curve indicated by the shaded regions.

We crushed various fruits, with a wide range of material characteristics, using a Material Testing System (MTS) universal testing machine (MTS Systems, Eden Prairie, Minnesota, USA) to detect clear differences in performance among fruit types. For our tests, we set a uniform loading rate of 30 mm/min (Crofts & Summers, 2014; Kolmann et al., 2015) with set displacements large enough to puncture the rind of the fruit but shallow enough to avoid collision and subsequent damage of the models (Figure 3). We tested sapote (*Pouteria - Sapotaceae*), cherimoya or soursop (*Annona - Annonaceae*), and mamoncillos (*Melicoccus - Sapindaceae*), all of which are fruits consumed by pacus and other characins in the flooded forest or domesticated cultivars of these same fruits (Goulding, 1980; Goulding et al., 1988; Kubitzki & Ziburski, 1994). We also included cherries (*Prunus - Rosaceae*) in our analysis, as a material analogue to palm fruit (*Astrocaryum* - Arecaceae), which pacus may encounter in their native habitat as well. Finally, we included kiwis (*Actinidia* - Actinidiaceae) and macadamia nuts (*Macadamia* - Proteaceae) to provide a further breadth of material characteristics.

We measured three performance metrics that correspond to ecologically-relevant aspects of food capture and processing: (1) load at puncture (Newtons - N), (2) work to maximum load (Newton meters-Nm) and (3) work to maximum depth (Nm). These metrics correspond to (1) the ability of a dentition to first indent and puncture fruit rind, then (2) pulverize pulp tissue, and finally (3) the work required to drive the tooth through the pulp tissue, which corresponds to the toughness of the fruit (Lucas; 2004; Hartstone-Rose et al., 2015).

### Gel visualization

To visualize changes in stress concentration during a ‘bite’ through a ductile material, we filmed polarized light through blocks of gelatine while running an MTS test as described above. We prepared ballistic gelatin in a 10% weight/volume ratio by combining KNOX unflavored gelatin with water, making sure to only use gels within 6 hours of their mixing (Knox gelatin, E.D. Smith Foods, Ltd.). To improve visualization, we melted the camera-facing side of each gelatine block with a small flame prior to testing and shone polarized light from the opposite side of the block while filming.

### Statistical analyses

Force and displacement data from individual trials were exported as .csv files from TestWorks software (MTS Systems, Eden Prairie, Minnesota, USA), reformatted using MATLAB (MathWorks, Natick Massachusetts, USA), and merged using R (project.org). We used a custom R script, *CrshR*, to read force-displacement curves into the RStudio GUI (https://github.com/CDonatelli/CrshR). For each force-displacement curve, this code automatically identifies maximum force (N) and maximum work (Nm), and allows users to assign a yield point, and calculates the force and work at that point. For the purposes of our study, we considered yield to be the point of the first significant inflection along the force-displacement slope and took this to indicate the point of initial puncture.

To contrast puncture performance among different serrasalmid species, we used two-way analysis of covariance (ANCOVA), with a given performance metric (force or work) as our dependent variable and species or diet as our independent variables. Our covariates were fruit or seed type and our various metrics of fruit/seed size. We used Tukey Honest Significant Difference (HSD) tests to evaluate significance of pairwise comparisons among species. This approach was repeated to compare fruit/seed materials, with the performance metric (force or work) as the dependent variable, fruit type as our independent variable, and fruit/seed size as a covariate.

Finally, we were interested in visualizing the puncture performance of a greater diversity of fruits, and used principal components analysis to construct a fruit/seed functional landscape. A p-value lesser than or equal to 0.05 was considered significant for all statistical tests. All analyses were performed in R (Version 2023.06.2+561).

## RESULTS

### Tooth indentation in fruits & nuts

Macadamia nuts required more force and work to puncture than kiwis and cherries, with cherries needing the least force and kiwis the least work. Macadamia nuts required significantly higher force to puncture (F = 92.27; p < 0.01) and work to puncture (F = 4.17; p = 0.01). Kiwis exhibited the highest work to max load, with macadamia nuts requiring less, and cherries requiring the least. ANOVA results also showed significantly higher work to max load (p < 0.01) for kiwis than cherries (p < 0.01) or macadamia nuts (F =13.17; p = 0.01).

When assessing these data via a PCA (Figure 4), all variables loaded positively on PC1. Force to puncture and work to puncture loaded negatively on PC2, with work to max load loading positively on PC2. Most fruits clustered to the left side of the plot, negatively on PC1 and spanning both sides of the origin on PC2 (Figure 4). This region is characterized by low to intermediate force at puncture, work to puncture, and work to max load. All fruits overlapped in the mechanical properties to some extent, but sapote mainly occupied the center of the biplot with one point in the main cluster of fruits; this could reflect our lower sample size for sapote (Figure 4). Mamoncillo, kiwi, and macadamia nuts exhibited the most variable mechanical properties in our data set and were also the most difficult fruits to indent, observationally. For kiwis in particular, this may have reflected varying stages of ripeness, although most data points for kiwi fell within the center left side of the biplot, similar to softer fruits like cherries and cherimoya (Figure 4).

**Figure 4.**
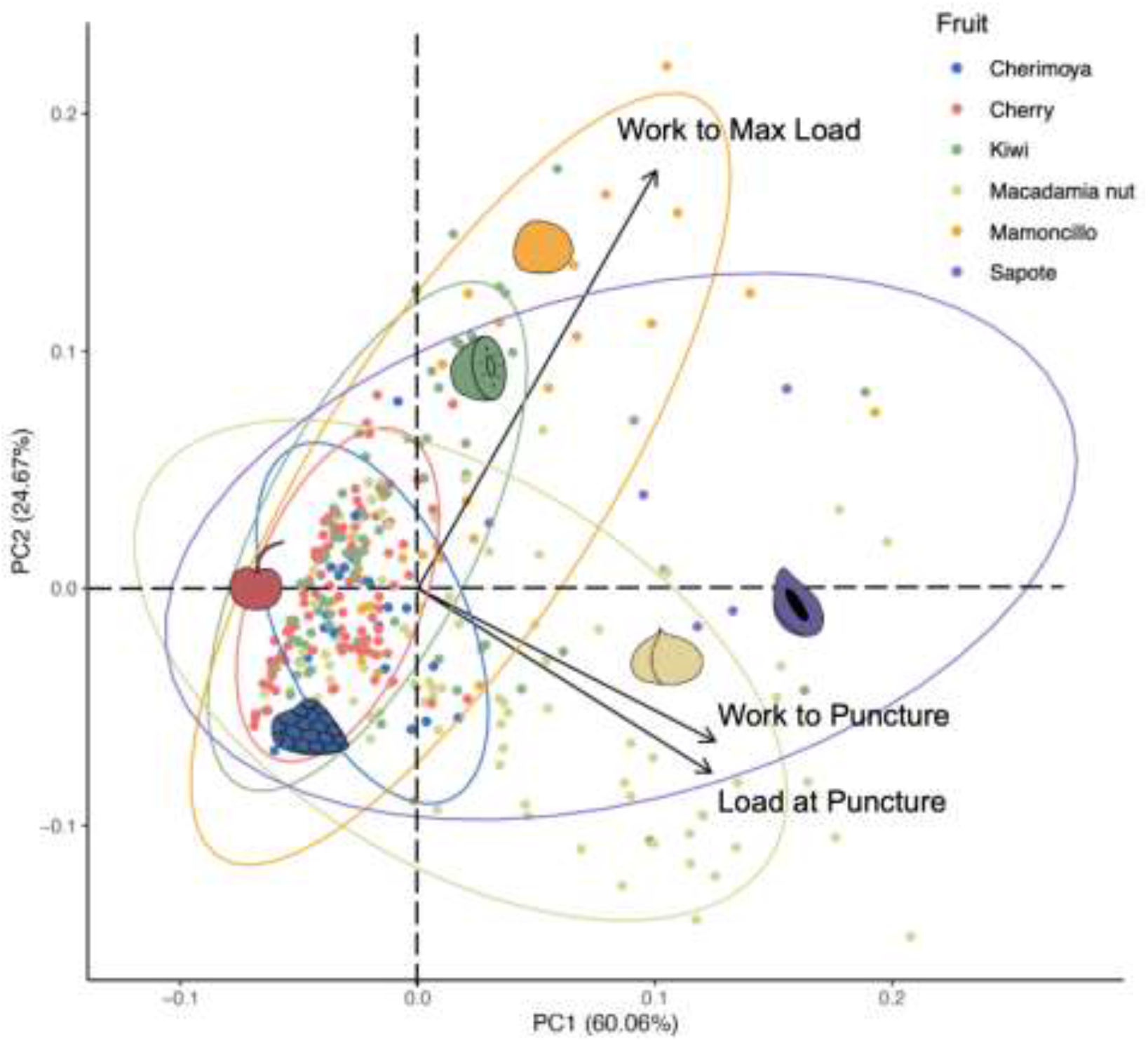
Principal components analysis describing the behaviors of different fruits. Vectors (arrows) represent the loadings of mechanical measures (load at puncture, work to puncture, work to max load) on the biplot. Cherimoya, mamoncillo, and sapote represent fruits most similar to the natural diet of pacus. Note that most fruits overlap in their mechanical behavior in the center left of the biplot.

### Tooth indentation performance among frugivorous and non-frugivorous serrasalmids

We also used the same three parameters as above (load at puncture, the work to puncture, and the work to max load) to contrast performance among the six serrasalmid species with the same three prey items. We focused on the differences between the frugivore, *Colossoma*, omnivores (*Prosomyleus, Myloplus, Acnodon, and Pristobrycon*), and *Serrasalmus*, a carnivore, in order to compare the puncture performance between species. In order to analyze across all six species we focused on puncture performance on cherries, kiwis, and macadamia nuts (Figure 5).

**Figure 5.**
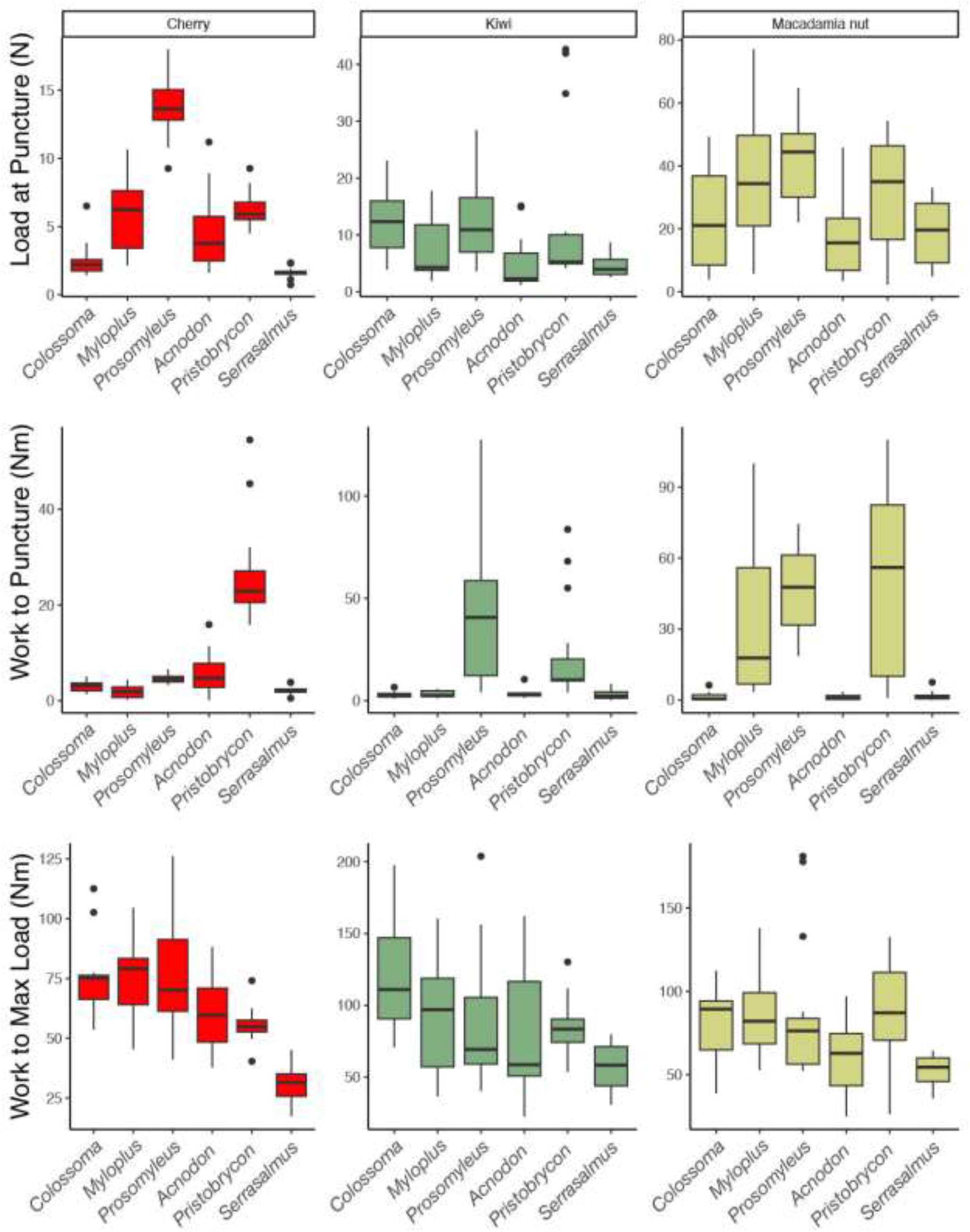
Boxplots of Force (N) and Work (Nm) done by serrasalmid dentitions on cherries (in red), kiwis (in green), and macadamia nuts (in yellow). (A) Load at Puncture (N), (B) Work to Puncture (Nm), (C) Work to Max Load (Nm). *Colossoma* (a pacu) and *Pristobrycon* (a piranha) are frugivores, *Serrasalmus* is a carnivore, while *Myloplus, Prosomyleus*, and *Acnodon* are omnivores.

For cherries, carnivorous *Serrasalmus* exhibited three times lower load at puncture with all species (*Prosomyleus, Myloplus, Acnodon*, and *Pristobrycon*) except the frugivore *Colossoma* which showed no significant difference. *Colossoma* required a significantly lower load at puncture than most of the omnivores, *Myloplus, Pristobrycon*, and *Prosomyleus. Prosomyleus* exhibited a significantly higher load at puncture than all other species (*Colossoma, Myloplus, Acnodon, Pristobrycon*, and *Serrasalmus*) (F = 70.22; p < 0.01). *Colossoma, Myloplus, Acnodon*, and *Serrasalmus* did not show significant difference in work to puncture but all exhibited significantly lower work to puncture when compared to *Prosomyleus* and *Pristobrycon*, which were not significantly different from one another (F = 25.77; p < 0.01). The piranhas, *Serrasalmus* and *Pristobrycon*, work to max load was significantly lower than that of the pacus (*Colossoma, Myloplus, Prosomyleus*, and *Acnodon*). While the work to max load was not significantly different between the pacus, *Pristobrycon* required significantly higher work to max load than *Serrasalmus*, which required half the work to max of *Colossoma* (F = 20.46; p ≤ 0.01) (Figure 5).

When testing with kiwi fruit, load at puncture was significantly lower for *Serrasalmus* relative to *Pristobrycon*, requiring more than half the load. Similarly, for load at puncture, *Prosomyleus* required double the force needed to indent fruit relative to *Serrasalmus*. Otherwise, there was no significant difference in load at puncture between the other species (F = 4.14; p < 0.04). For work to puncture the kiwi fruit, the omnivore *Prosomyleus* required significantly more work than all other species, with no other differences observed among species (F = 13.23; p ≤ 0.01). The frugivorous *Colossoma* required more than double the work to max load compared to the carnivore, *Serrasalmus* and almost double that of an omnivore *Acnodon*, and was significantly higher than both (F = 4.86; p ≤ 0.03) (Figure 5).

For models tested on macadamia nuts, *Prosomyleus* required significantly higher load at puncture when compared with *Colossoma, Serrasalmus*, or *Acnodon. Acnodon* also required a significantly lower load at puncture than *Myloplus* (F = 5.94; p ≤ 0.02). In comparing some of the more omnivorous species, *Prosomyleus* showed significantly higher work to puncture relative to *Acnodon* or *Colossoma*; *Myloplus* showed significantly greater work to puncture than *Acnodon* and *Colossoma*; and *Acnodon* was significantly lower than *Pristobrycon* as well. Both the frugivore (*Colossoma*) and the carnivore (*Serrasalmus*), required significantly less work to puncture than *Pristobrycon*, the omnivorous piranha (F = 18.81; p < 0.01). *Serrasalmus* required significantly lower work to max load than the omnivores, *Myloplus* and *Prosomyleus*, and the other piranha *Pristobrycon* (F = 4.62; p ≤ 0.02) (Figure 5).

### Gel visualization of how teeth propagate stress

There was a standard progression of events for all gell visualization tests: Load was applied and bands would appear, indicating concentrations of stress. The gel would fracture at failure, releasing stored energy and decreasing overall stress concentration in the block. Finally, tooth models were withdrawn from the gel block. There were clear differences in bite stress distributions between the piranha and pacu dentitions (Figure 6).

**Figure 6.**
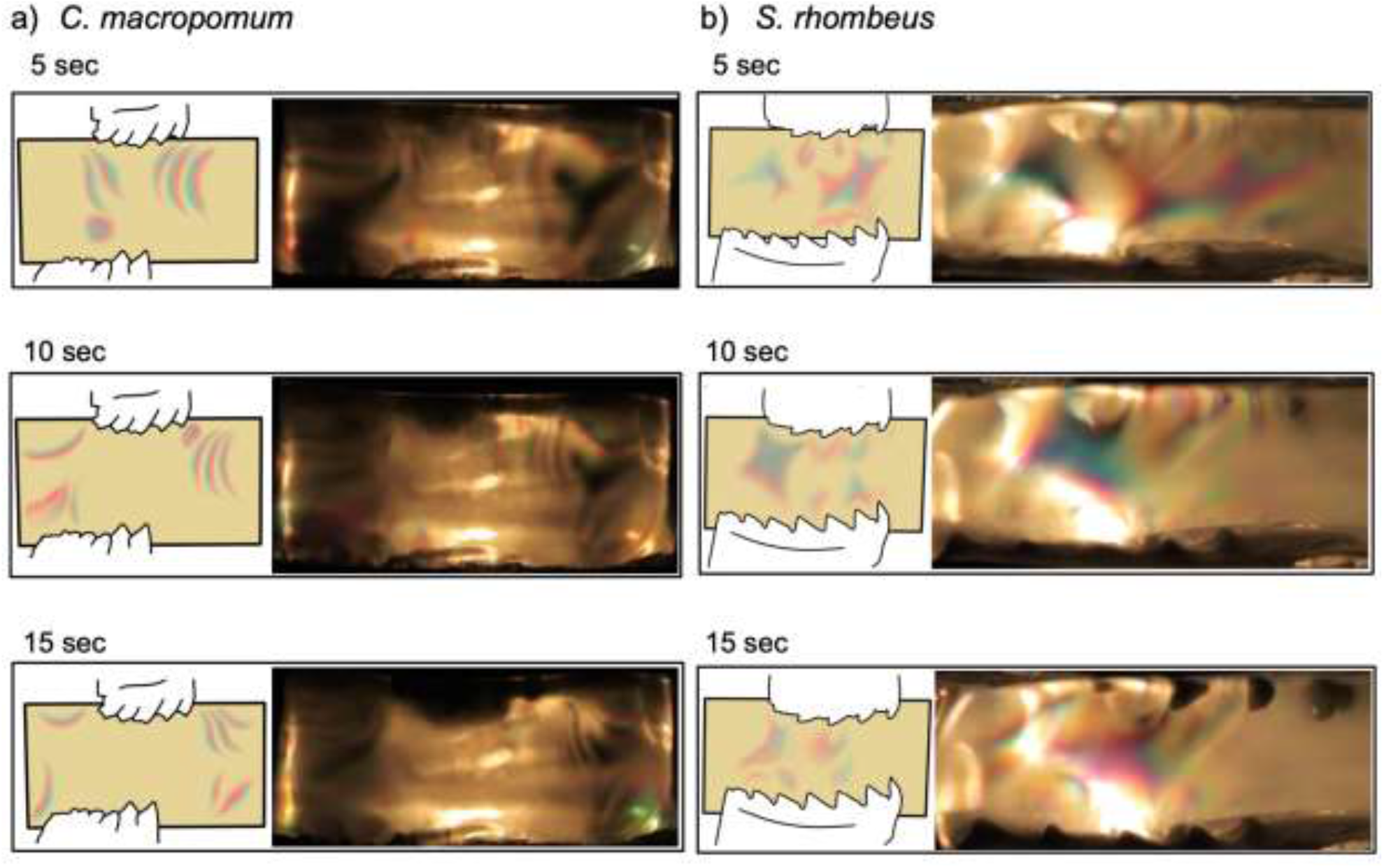
Gel visualization of (A) pacu and (B) piranha tooth indentation with corresponding schematics of stress distributions as visualized from these images. Colored portions indicate stress, non-colored portions indicate evenly distributed stress.

The dentition of pacus (*Colossoma, Prosomyleus, Acnodon*, and *Myloplus*) applied areas of high stress spread more evenly through the gel for the duration of loading. For the *Colossoma* (specialist frugivore) trial, lines of stress appeared from the dentition as a whole and were (very quickly) evenly distributed. Upon gel failure, we observed a release of stress by the bottom dentition, and on the removal of the dentition, the isochromatic fringes appear in the opposite direction than they did during puncture.

In both piranhas, *Serrasalmus* and *Pristobrycon*, stress was increasingly concentrated around the individual teeth as load was applied. Only after prey failure, and during withdrawal, did stress spread more evenly across the dentitions. As the *Serrasalmus* dentition entered the gel there was an immediate and intense stress produced in between individual teeth (Figure 6). During load trials, *Pristobrycon* exhibited rings of stress around the points of the teeth upon puncture, and a band of high stress appeared in the middle of the gel, towards the end of the modeled bite.

The *Acnodon* dentition produces an initial high stress at the isochromatic fringes, which moves laterally as the load increases, following the shape of the multicuspid teeth. The fourth and fifth teeth in the labial row of *Acnodon* are oriented in opposite directions; in these areas stress remained high throughout the applied load (Figure 6). During the gel trials of *Myloplus*, another omnivore, we observed greater birefringence at the rostral/caudal edges of the models rather than below but, as the gel failed and cracked, stress was quickly released. This was observed as the isochromatic rings condensed and then quickly expanded with minimal distance between each ring, suggesting that stress and energy both decreased. A similar trend in stress and strain was observed in the dentition trials of the third omnivorous species, *Prosomyleus*.

## DISCUSSION

### Low Force, High Work… and diversity in the performance of frugivorous fish dentitions

Vertebrates that lack nimble digits, such as frugivorous bats and fishes, would appear to face a distinct challenge in dismantling and consuming fruits when compared to frugivorous primates, using teeth rather than digits to manipulate food as well as process it. Teeth must be sharp enough to puncture tough fruit rinds to effectively maintain a grip (i.e., require relatively low initial force and work), an action that is especially challenging for gape-limited fishes (Alexander, 1964) like *Colossoma* that process fruits exceeding the size of their mouth. *Colossoma* and other pacus are known to stake out fruiting trees in groups, waiting for fruit to fall (Goulding, 1980) - so a secure grip on fruit will be paramount as fish contend with schoolmates trying to abscond with their meal.

Beyond the point of the initial puncture and grip, fish are faced with the additional challenge of processing the fruit. While the relatively soft tissues of these fruits (like pulp) deform easily, potentially impeding rack propagation (Lucas, 1979; Lucas & Luke, 1984), the assembly of teeth in the jaw creates notches, trapping the soft tissues and concentrating applied force (Anderson, 2009; Anderson and LaBarbera, 2008). In carnivorous piranhas, like *Serrasalmus*, teeth are bladed, so this concentrated force is applied over the relatively small surface area of the tooth blade, overall reducing the stress needed to cut through tough flesh force (Anderson, 2009; Anderson and LaBarbera, 2008). Frugivorous fishes face a less obvious challenge; they need to access the sugars within the soft tissue of the fruit, rather than simply slice through it. For this to happen, tooth surfaces adjacent to the cusps need to be blunt enough to burst cell walls in the pulp (i.e., generate high work).

In line with our predicted model for frugivory, dentition of the frugivorous *Colossoma* required relatively low force and work to initially puncture fruits (cherry and kiwi), and a higher amount of work to reach maximum force (Figure 5). Moreover, carnivorous piranhas, like *Serrasalmus*, with teeth that required both low force and low work or process food items, were most readily distinguishable in performance from frugivores among the serrasalmids (Figure 5).

These findings further supported the observations from our series of gel visualization experiments. The gel birefringence experiments revealed that pacu dentitions exert a more even stress distribution across prey than the dentitions of the more carnivorous piranhas, which generate high stress only in localized regions. We found that the central cusp of piranha teeth concentrates high stresses over a narrow area of the gel, increasing the rate at which localized stresses accumulate and facilitating penetration into the material. After this initial high stress at puncture there is a release of stress and embedded piranha teeth ‘slide’ deeper into the gel with little resistance. The concentration of stresses around the individual teeth of *Serrasalmus*, the piranha, suggests a priority for speed through the material and removal of the bitten portion of the prey item.

Not all piranhas have carnivorous diets, some may be omnivorous (Nico & Taphorn, 1988; Prudente et al., 2016) or herbivorous (Kolmann et al., 2024). High performance feeding, usually through generation of higher bite forces, can serve to expand niche breadth in other vertebrates (Hernandez & Motta, 1997; Verwaijen et al., 2002), which would seem beneficial to omnivores and specialists alike. Piranha teeth are typically blade-like, adapted to puncture prey with low resistance, engage efficiently and stab deeply into prey tissues. *Pristobrycon maculipinnis*, a poorly-known piranha from the Venezuelan llanos, and what little is known about the ecology suggests that it, at least in some part, is frugivorous (Nico, pers comm; Nico, 1991). Our work here supports this proposed diet, as the *Pristobrycon* dentition required higher force and much higher work than *Serrasalmus* to process the different fruits (Figure 5). This “high force, high work” pattern is consistent with the “mortar and pestle” model (Figure 2), of teeth adapted for crushing ‘hard’ food, more like hard seeds than fleshy fruits. Similarly, one of the omnivores included in this study, *Prosomyleus*, also had data consistent with the “mortar and pestle” model requiring comparably higher forces to puncture cherries and macadamia nuts. Which suggests that *Prosomyleus*, like *Pristobrycon*, may be more adapted to durophagy (and granivory). We suggest fishes like *Prosomyleus* and *Pristobrycon* may be a second kind of frugivore, one which targets the seeds, rather than the flesh, of fruits. In a hypothetical spectrum from seed predator to seed disperser, where a frugivore obliterates all ingested seeds in the former and excretes most seeds intact (and alive) in the latter, we propose that “mortar and pestle” dentitions are associated with seed predation.

### Diversity of fruit material properties & frugivore dentitions

Variation in fruit ripeness significantly impacts the performance of dentitions during biting, suggesting the tooth morphology and cusp arrangement is optimized for a particular ripeness. For example, an unripe kiwi is uniformly hard, while ripe kiwis have a tough rind and soft inner flesh. The stiff flesh of an unripe fruit causes more difficulty in puncturing the fruit initially but makes propagating a crack through the fruit and breaking it apart easier (Lucas & Corlett, 1991). For frugivores, crushing a tough, unripe fruit’s hard flesh requires much more work (Hartstone-Rose et al., 2015) and provides presumably less caloric payout than consuming softer flesh in ripe fruits (Lambert, 1999), particularly when secondary compounds in unripe fruit can harm frugivores. While excellent at cutting through fruit tissues and driving cracks through stiff materials (like seeds), the blade-like teeth of piranhas function less well at pulverizing fruit cells.This is perhaps one additional piece of evidence that frugivory in omnivorous piranhas may be more driven by seed predation than a focus on fruit. Teeth in omnivorous piranhas can slice cleanly through pulp, thereby accessing and then fracturing the seeds inside. This mechanical interpretation aligns with some natural history observations, where seeds ingested by piranhas are frequently macerated (in ‘*Serrasalmus’ striolatus*, Goulding, 1980; *S. goulding*, Prudente et al., 2016; in *Pygopristis*, Kolmann et al., 2024). Moreover, the same pattern of finding sharper tooth cusps in frugivores that also engage in seed predation is mirrored in prosimians like lemurs (Yamashita, 1996; Overdorff & Strait, 1998).

One could question the ecological relevance of our fruit sample, given that pacus and piranhas, outside the domestic aquarium, generally do not dine on cherries. However, our methods reveal that the properties of more common commercially-available fruits like cherries and kiwis overlapped with fruits like cherimoya and mamoncillo (Figure 3). These less commercially-available fruits belong to the same families (Annonaceae and Sapindaceae, respectively) that pacus and piranhas consume in nature (Goulding, 1980). In fact, cherimoya (*Annona cherimola*) is a domesticated cultivar (i.e., in the same genus) of one of the fruits that pacus consume in tropical lowlands (e.g., *Annona muricata*; Anderson et al., 2009). Unfortunately, it is exceedingly difficult to access the fruits and seeds typically found in the diets of frugivorous fishes and other frugivorous vertebrates in general. However, this study and others, which use non-natural fruits for mechanical and behavioral assays (e.g., Dumont, 1999; Hartstone-Rose et al., 2015), can still provide insight into the form-function relationships between frugivores and their food, at least in general.

### Convergence in frugivorous vertebrate dentitions

The dentition of *Colossoma* and other frugivorous serrasalmids also have morphological characteristics in common with other vertebrate frugivores like bats, namely: (1) ‘cookie-cutter’ like occlusion of upper and lower teeth (Freeman, 1992), (2) sharp and near continuous labial cutting surfaces on the dentition (Freeman 1992), and (3) plier-like gripping occlusion (relative to more scissor-like occlusion in animalivorous relatives; Freeman, 1988). The former two dental traits may be useful in removing rind from fruit without damaging soft pulp prior to prey processing or ‘juicing’ (Freeman, 1992). Previous studies on serrasalmid feeding morphologies have found gripping or crushing dental occlusion in herbivores in general (Huby et al., 2019) and frugivores specifically (Kolmann et al., 2024).

Frugivorous pacus also appear to share the following mandibular morphologies in common with terrestrial frugivores (Freeman, 1988, 1992; Dumont 1997, 1999, 2004; Vogel et al., 2014): a U-shaped lower jaw with strongly interdigitated or fused medial symphyses (Kolmann, pers comm; Figure 1), and a semi-prehensile lip (Cohen et al., 2023). A strongly interdigitated or fused mandibular symphysis, as in serrasalmids and mammals, is beneficial here for effectively transmitting unilateral muscle forces across the symphysis and resisting anterior reaction forces (Hylander, 1979; Freeman, 1992). Finally, like the tongue and lips in frugivorous primates (Lambert, 1999), fruit-feeding pacus are thought to use a semi-prehensile lip to help in manipulating and repositioning food (Cohen et al., 2023).

## ACKNOWLEDGEMENTS

The authors dedicate the manuscript to Michael Goulding for writing his influential book, ‘*The Fishes and the Forest*,’ the inspiration for this work. Hernán López-Fernández and Ramon Nagesan (University of Michigan Museum of Zoology) facilitated CT scanning of the *Pristobrycon maculipinnis* specimen for this study. JR thanks the Steven and Ruth Wainwright Endowment. Funding was provided by NSF-PRFB 1712015 and University of Louisville start-up funds to MAK. CD was supported by Friday Harbor Labs. KC was supported by the Steven Wainwright Foundation. SC was supported by College of the Holy Cross start-up funds. APS was supported by the Seaver Institute and the National Science Foundation.

## AUTHOR CONTRIBUTIONS

MAK, SC, and JR conceived and designed the study. JR collected the data. KEC and JR collected and analyzed gel visualization data. JR analyzed the data. CD wrote custom MatLab and R scripts to collate and process loading frame data. JR and MAK drafted the initial version of the manuscript, and all authors contributed to final versions of the manuscript.

## DATA ACCESSIBILITY

All microCT scan data, indentation (force/load, displacement) data, and gel visualization videos are deposited in a Dryad digital repository [doi: *tbd*].

## SUPPLEMENTARY TABLES

**S1a.** Kiwi, Load at puncture. Analysis of Variance (ANOVA) results for indentation tests. Significance was considered with an alpha greater than or equal to 0.05.

|  | diff | lwr | upr | adj. p-value |
| --- | --- | --- | --- | --- |
| *Myloplus-Colossoma* | -4.881 | -12.953 | 3.191 | 0.495 |
| *Prosomyleus-Colossoma* | 0.498 | -7.574 | 8.570 | 0.999 |
| *Acnodon-Colossoma* | -7.378 | -15.449 | 0.694 | 0.093 |
| *Pristobrycon-Colossoma* | 0.602 | -7.470 | 8.674 | 0.999 |
| *Serrasalmus-Colossoma* | -7.750 | -15.822 | 0.322 | 0.067 |
| *Prosomyleus-Myloplus* | 5.379 | -2.693 | 13.451 | 0.384 |
| *Acnodon-Myloplus* | -2.497 | -10.569 | 5.575 | 0.945 |
| *Pristobrycon-Myloplus* | 5.483 | -2.589 | 13.555 | 0.362 |
| *Serrasalmus-Myloplus* | -2.869 | -10.941 | 5.203 | 0.904 |
| *Acnodon-Prosomyleus* | -7.876 | -15.947 | 0.196 | 0.060 |
| *Pristobrycon-Prosomyleus* | 0.104 | -7.968 | 8.176 | 1.000 |
| *Serrasalmus-Prosomyleus* | -8.248 | -16.320 | -0.176 | 0.042 |
| *Pristobrycon-Acnodon* | 7.980 | -0.092 | 16.052 | 0.054 |
| *Serrasalmus-Acnodon* | -0.372 | -8.444 | 7.700 | 0.999 |
| *Serrasalmus-Pristobrycon* | -8.352 | -16.424 | -0.280 | 0.038 |

**S1b.** Kiwi, Work to puncture. Analysis of Variance (ANOVA) results for indentation tests. Significance was considered with an alpha greater than or equal to 0.05.

|  | diff | lwr | upr | adj. p-value |
| --- | --- | --- | --- | --- |
| *Myloplus-Colossoma* | 0.328 | -19.535 | 20.190 | 1.000 |
| *Prosomyleus-Colossoma* | 42.613 | 22.751 | 62.475 | 0.000 |
| *Acnodon-Colossoma* | 0.685 | -19.177 | 20.548 | 0.999 |
| *Pristobrycon-Colossoma* | 19.676 | -0.186 | 39.539 | 0.054 |
| *Serrasalmus-Colossoma* | 0.080 | -19.783 | 19.942 | 1.000 |
| *Prosomyleus-Myloplus* | 42.285 | 22.423 | 62.148 | 0.000 |
| *Acnodon-Myloplus* | 0.358 | -19.505 | 20.220 | 1.000 |
| *Pristobrycon-Myloplus* | 19.349 | -0.514 | 39.211 | 0.061 |
| *Serrasalmus-Myloplus* | -0.248 | -20.110 | 19.614 | 1.000 |
| *Acnodon-Prosomyleus* | -41.928 | -61.790 | -22.065 | 0.000 |
| *Pristobrycon-Prosomyleus* | -22.937 | -42.799 | -3.074 | 0.014 |
| *Serrasalmus-Prosomyleus* | -42.533 | -62.396 | -22.671 | 0.000 |
| *Pristobrycon-Acnodon* | 18.991 | -0.871 | 38.853 | 0.069 |
| *Serrasalmus-Acnodon* | -0.606 | -20.468 | 19.257 | 1.000 |
| *Serrasalmus-Pristobrycon* | -19.597 | -39.459 | 0.266 | 0.055 |

**S1c.**
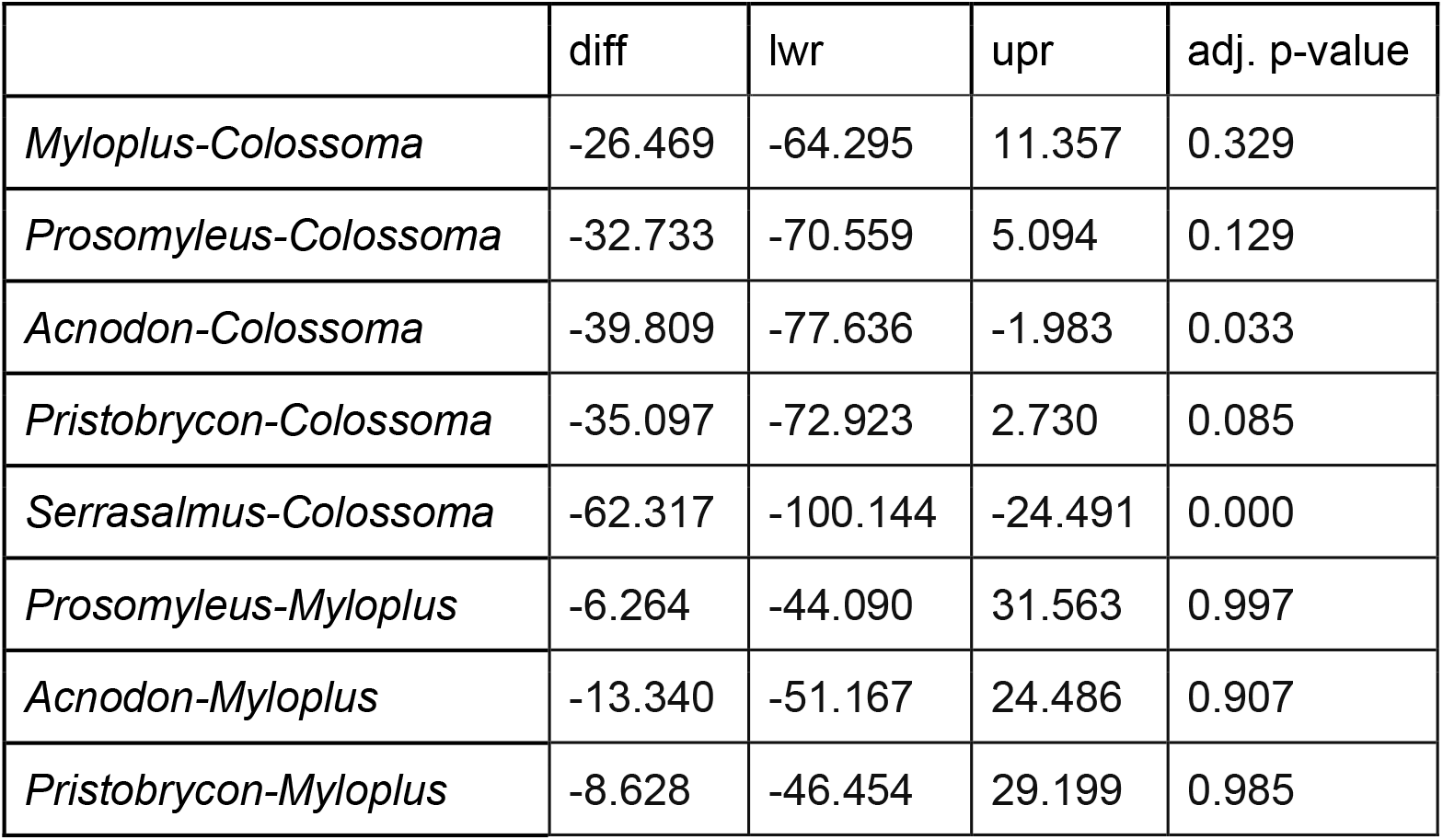
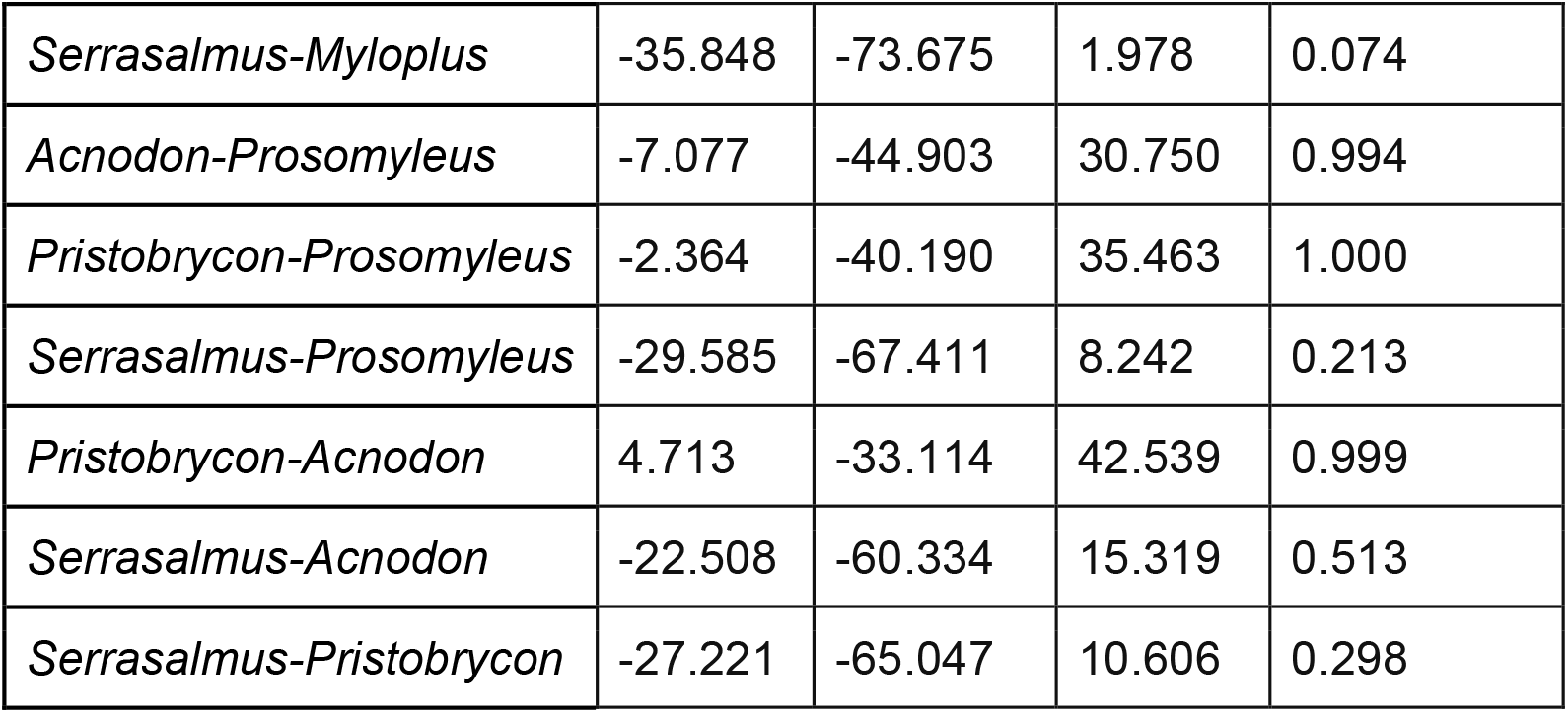
Kiwi, Work to max load. Analysis of Variance (ANOVA) results for indentation tests. Significance was considered with an alpha greater than or equal to 0.05.

**S2a.** Macadamia nuts, Load at puncture. Analysis of Variance (ANOVA) results for indentation tests. Significance was considered with an alpha greater than or equal to 0.05.

|  | diff | lwr | upr | adj. p-value |
| --- | --- | --- | --- | --- |
| *Myloplus-Colossoma* | 12.431 | -4.035 | 28.896 | 0.248 |
| *Prosomyleus-Colossoma* | 18.912 | 2.447 | 35.377 | 0.015 |
| *Acnodon-Colossoma* | -5.229 | -21.694 | 11.237 | 0.939 |
| *Pristobrycon-Colossoma* | 8.709 | -7.757 | 25.174 | 0.638 |
| *Serrasalmus-Colossoma* | -3.998 | -20.464 | 12.467 | 0.980 |
| *Prosomyleus-Myloplus* | 6.481 | -9.984 | 22.947 | 0.860 |
| *Acnodon-Myloplus* | -17.659 | -34.125 | -1.194 | 0.028 |
| *Pristobrycon-Myloplus* | -3.722 | -20.187 | 12.743 | 0.986 |
| *Serrasalmus-Myloplus* | -16.429 | -32.894 | 0.036 | 0.051 |
| *Acnodon-Prosomyleus* | -24.141 | -40.606 | -7.675 | 0.001 |
| *Pristobrycon-Prosomyleus* | -10.203 | -26.668 | 6.262 | 0.467 |
| *Serrasalmus-Prosomyleus* | -22.910 | -39.376 | -6.445 | 0.002 |
| *Pristobrycon-Acnodon* | 13.937 | -2.528 | 30.403 | 0.145 |
| *Serrasalmus-Acnodon* | 1.230 | -15.235 | 17.696 | 0.999 |
| *Serrasalmus-Pristobrycon* | -12.707 | -29.172 | 3.758 | 0.226 |

**S2b.**
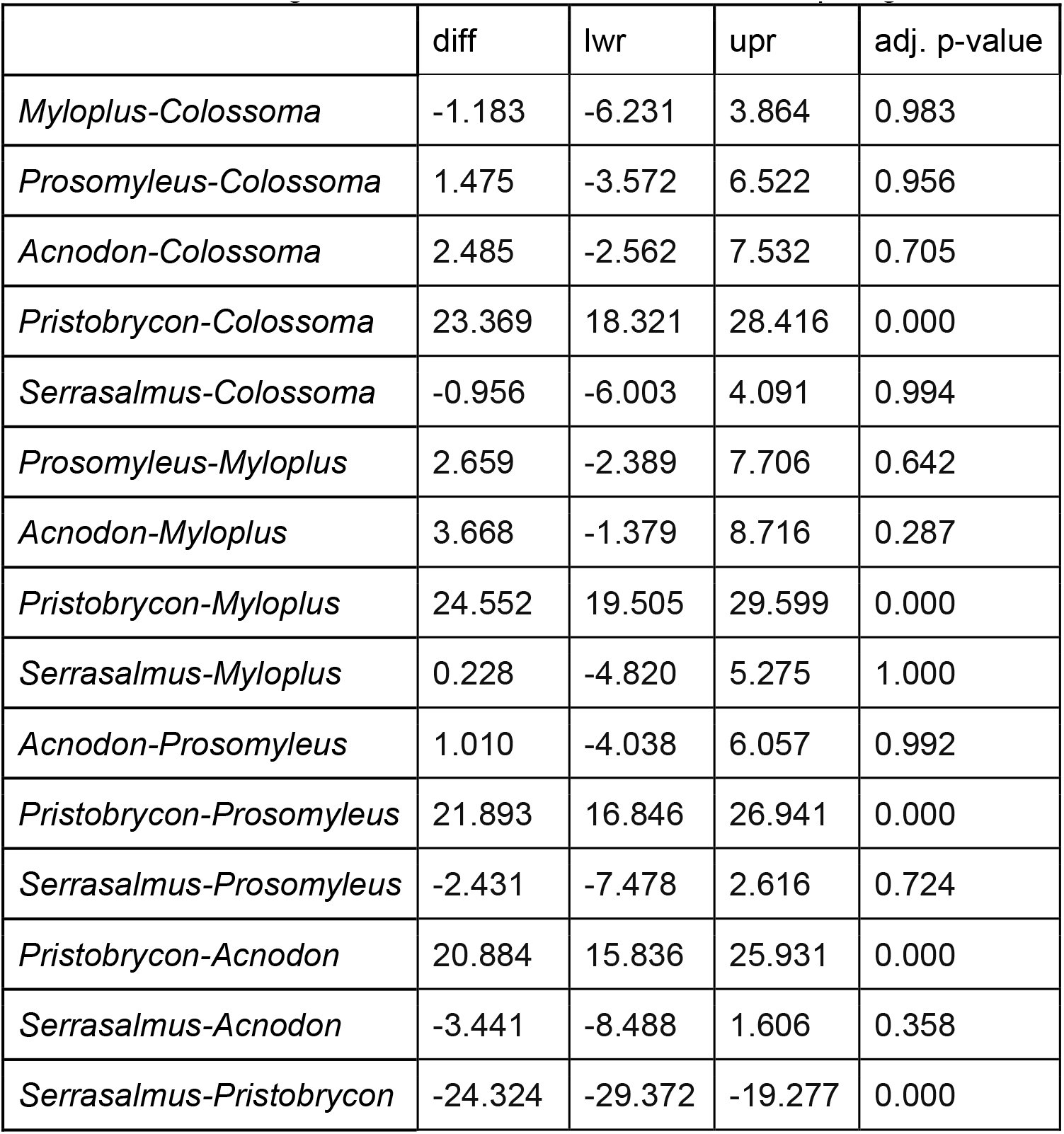
Macadamia nuts, Work to puncture. Analysis of Variance (ANOVA) results for indentation tests. Significance was considered with an alpha greater than or equal to 0.05.

**S2c.**
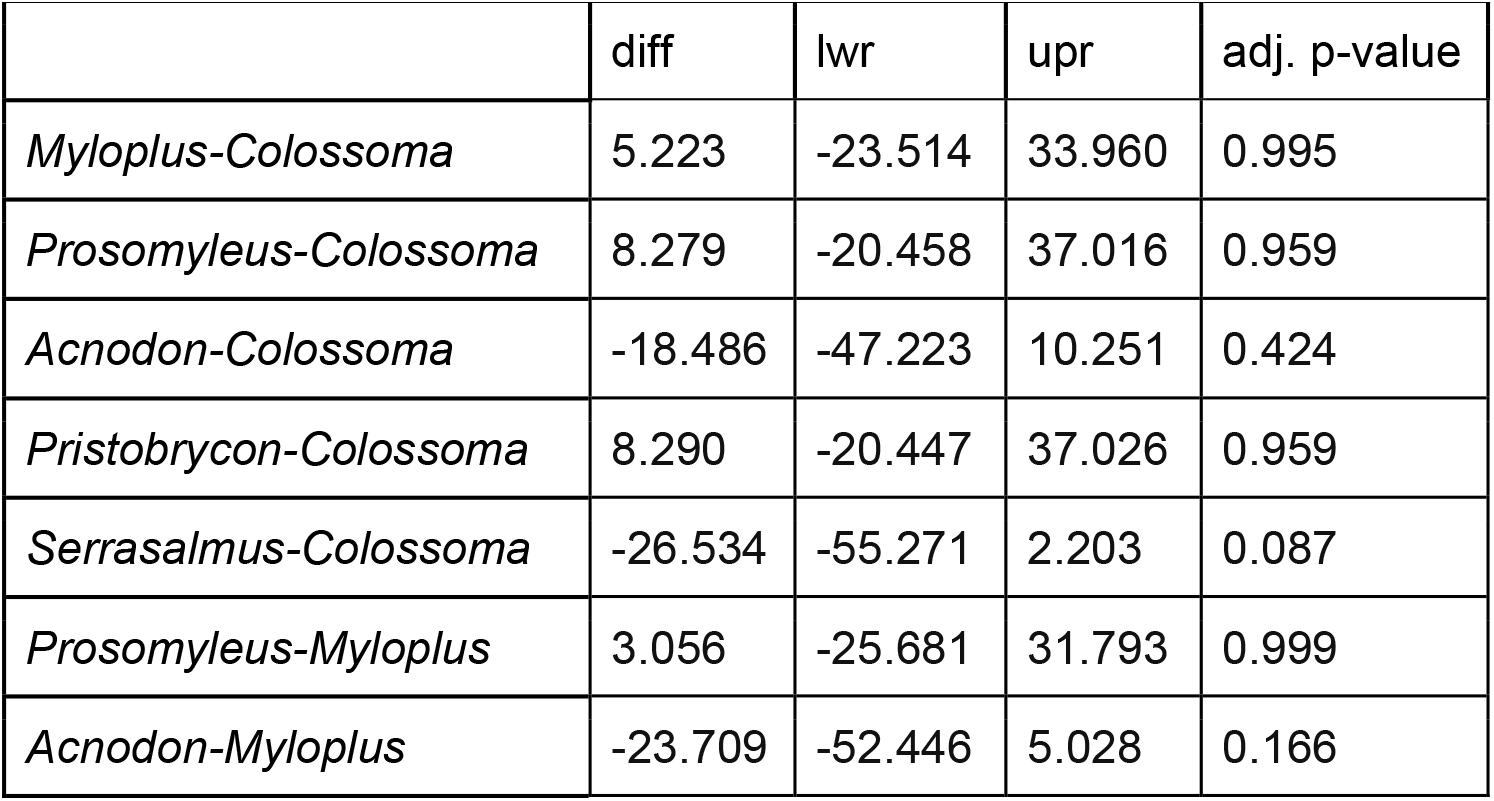
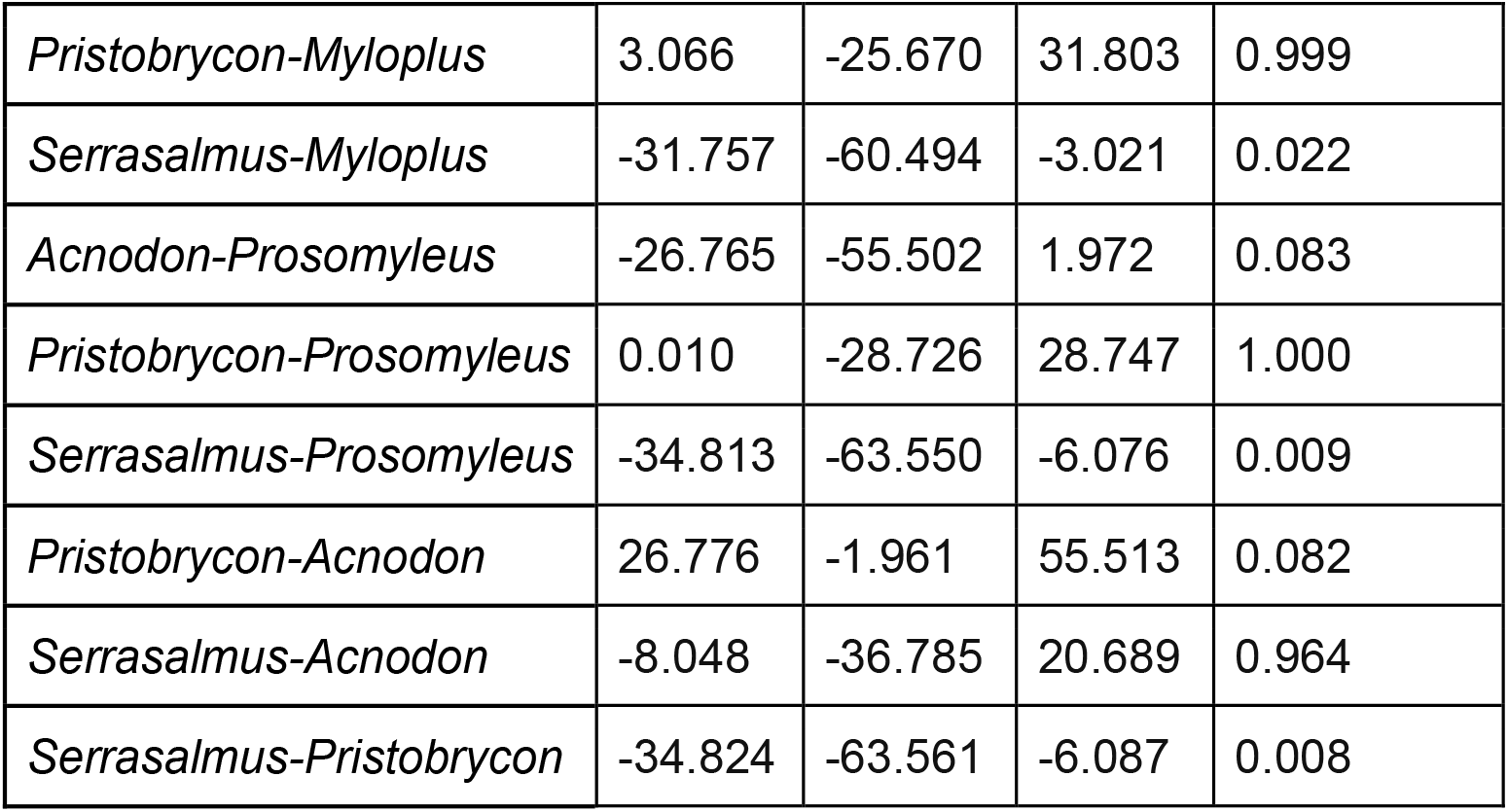
Macadamia nuts, Work to max load. Analysis of Variance (ANOVA) results for indentation tests. Significance was considered with an alpha greater than or equal to 0.05.

|  | diff | lwr | upr | adj. p-value |
| --- | --- | --- | --- | --- |
| *Myloplus-Colossoma* | 5.223 | -23.514 | 33.960 | 0.995 |
| *Prosomyleus-Colossoma* | 8.279 | -20.458 | 37.016 | 0.959 |
| *Acnodon-Colossoma* | -18.486 | -47.223 | 10.251 | 0.424 |
| *Pristobrycon-Colossoma* | 8.290 | -20.447 | 37.026 | 0.959 |
| *Serrasalmus-Colossoma* | -26.534 | -55.271 | 2.203 | 0.087 |
| *Prosomyleus-Myloplus* | 3.056 | -25.681 | 31.793 | 0.999 |
| *Acnodon-Myloplus* | -23.709 | -52.446 | 5.028 | 0.166 |
| *Pristobrycon-Myloplus* | 3.066 | -25.670 | 31.803 | 0.999 |
| *Serrasalmus-Myloplus* | -31.757 | -60.494 | -3.021 | 0.022 |
| *Acnodon-Prosomyleus* | -26.765 | -55.502 | 1.972 | 0.083 |
| *Pristobrycon-Prosomyleus* | 0.010 | -28.726 | 28.747 | 1.000 |
| *Serrasalmus-Prosomyleus* | -34.813 | -63.550 | -6.076 | 0.009 |
| *Pristobrycon-Acnodon* | 26.776 | -1.961 | 55.513 | 0.082 |
| *Serrasalmus-Acnodon* | -8.048 | -36.785 | 20.689 | 0.964 |
| *Serrasalmus-Pristobrycon* | -34.824 | -63.561 | -6.087 | 0.008 |

## REFERENCES

Alexander, R.M., 1964. Adaptation in the skulls and cranial muscles of South American characinoid fish. Zoological Journal of the Linnean Society, 45(305), pp.169–190.

Anderson, J.T., Rojas, J.S. and Flecker, A.S., 2009. High-quality seed dispersal by fruit-eating fishes in Amazonian floodplain habitats. Oecologia, 161(2), pp.279–290.

Anderson, J.T., Nuttle, T., Saldaña Rojas, J.S., Pendergast, T.H. and Flecker, A.S., 2011. Extremely long-distance seed dispersal by an overfished Amazonian frugivore. Proceedings of the Royal Society B: Biological Sciences, 278(1723), pp.3329–3335.

Andrade, M.C., Giarrizzo, T. and Jégu, M., 2013. Tometes camunani (Characiformes: Serrasalmidae), a new species of phytophagous fish from the Guiana Shield, rio Trombetas basin, Brazil. Neotropical Ichthyology, 11(2), pp.297–306.

Andrade, M.C., Jégu, M. and Giarrizzo, T., 2016a. Tometes kranponhah and Tometes ancylorhynchus (Characiformes: Serrasalmidae), two new phytophagous serrasalmids, and the first Tometes species described from the Brazilian Shield. Journal of Fish Biology, 89(1), pp.467–494.

Andrade, M.C., M. Jégu and T. Giarrizzo. 2016b. A new large species of Myloplus (Characiformes, Serrasalmidae) from the Rio Madeira basin, Brazil. ZooKeys 57: 153–167.

Andrade, M.C., M. Jégu, P.A. Buckup and A.L. Netto-Ferreira. 2018. A new Myleus species (Characiformes: Serrasalmidae) from the Rio Tapajós basin, Brazil.Journal of Fish Biology.

Andrade, M.C., Fitzgerald, D.B., Winemiller, K.O., Barbosa, P.S. and Giarrizzo, T., 2019a. Trophic niche segregation among herbivorous serrasalmids from rapids of the lower Xingu River, Brazilian Amazon. Hydrobiologia, 829(1), pp.265–280.

Araujo, J.M., Correa, S.B., Anderson, J. and Penha, J., 2020. Fruit preferences by fishes in a Neotropical floodplain. Biotropica, 52(6), pp.1131–1141.

Araujo, J.M., Correa, S.B., Penha, J., Anderson, J. and Traveset, A., 2021. Implications of overfishing of frugivorous fishes for cryptic function loss in a Neotropical floodplain. Journal of Applied Ecology, 58(7), pp.1499–1510.

Augspurger, C.K., 1983. Seed dispersal of the tropical tree, Platypodium elegans, and the escape of its seedlings from fungal pathogens. The Journal of Ecology, pp.759–771.

Betancur-R, R., Arcila, D., Vari, R.P., Hughes, L.C., Oliveira, C., Sabaj, M.H. and Ortí, G., 2019. Phylogenomic incongruence, hypothesis testing, and taxonomic sampling: The monophyly of characiform fishes. Evolution, 73(2), pp.329–345.

Boujard, T., Sabatier, D., Rojas-Beltran, R., Prevost, M.F. and Renno, J.F., 1990. The food habits of three allochthonous feeding characoids in French Guiana. Revue d’Ecologie, Terre et Vie, 45(3), pp.247–258.

Burns, M.D., 2021. Adaptation to herbivory and detritivory drives the convergent evolution of large abdominal cavities in a diverse freshwater fish radiation (Otophysi: Characiformes). Evolution, 75(3), pp.688–705.

Burns, M.D. and Sidlauskas, B.L., 2019. Ancient and contingent body shape diversification in a hyperdiverse continental fish radiation. Evolution, 73(3), pp.569–587.

Buser, T.J., Boyd, O.F., Cortés, Á., Donatelli, C.M., Kolmann, M.A., Luparell, J.L., Pfeiffenberger, J.A., Sidlauskas, B.L. and Summers, A.P., 2020. The natural historian’s guide to the CT galaxy: step-by-step instructions for preparing and analyzing computed tomographic (CT) data using cross-platform, open access software. Integrative Organismal Biology, 2(1), p.obaa009.

Chivers, D.J., Andrews, P., Preuschoft, H., Bilsborough, A. and Wood, B.A., 1984. Food acquisition and processing in primates: concluding discussion. In Food acquisition and processing in Primates (pp. 545–556). Boston, MA: Springer US.

Cignoni, M. Callieri, M. Corsini, M. Dellepiane, F. Ganovelli, G. Ranzuglia, MeshLab: an Open-Source Mesh Processing Tool, Sixth Eurographics Italian Chapter Conference, page 129–136, 2008

Cione, A.L., Dahdul, W.M., Lundberg, J.G. and Machado-Allison, A., 2009. Megapiranha paranensis, a new genus and species of Serrasalmidae (Characiformes, Teleostei) from the upper Miocene of Argentina. Journal of Vertebrate Paleontology, 29(2), pp.350–358.

Cohen, K.C., Lucanus, O., Summers, A.P., Kolmann, M.A. 2023. Lip service: histological phenotypes correlate with diet and feeding ecology in herbivorous pacus. Anatomical Record. 306 (2), 326–342.

Correa, S.B., Winemiller, K.O., Lopez-Fernandez, H. and Galetti, M., 2007. Evolutionary perspectives on seed consumption and dispersal by fishes. Bioscience. 57(9), pp.748–756.

Correa, S.B. and Winemiller, K.O., 2014. Niche partitioning among frugivorous fishes in response to fluctuating resources in the Amazonian floodplain forest. Ecology. 95(1), pp.210–224.

Correa, S.B., Costa-Pereira, R., Fleming, T., Goulding, M. and Anderson, J.T., 2015. Neotropical fish–fruit interactions: eco-evolutionary dynamics and conservation. Biological Reviews. 90(4), pp.1263–1278.

Correa, S.B., Arujo, J.K., Penha, J., Nunes da Cunha, C., Bobier, K.E. and Anderson, J.T., 2016. Stability and generalization in seed dispersal networks: a case study of frugivorous fish in Neotropical wetlands. Proceedings of the Royal Society B: Biological Sciences. 283(1837), p.20161267.

Correa, S.B., de Oliveira, P.C., Nunes da Cunha, C., Penha, J. and Anderson, J.T., 2018. Water and fish select for fleshy fruits in tropical wetland forests. Biotropica. 50(2): 312–318.

Costa-Pereira, R., Correa, S.B. and Galetti, M., 2018. Fishing-down within populations harms seed dispersal mutualism. Biotropica, 50(2), pp.319–325.

Crofts, S. B., Smith, S. M. and Anderson, P. S. L. (2020). Beyond Description: The Many Facets of Dental Biomechanics. Integr. Comp. Biol. 60.

Dary EP, Ferreira E, Zuanon J, Röpke CP. 2017. Diet and trophic structure of the fish assemblage in the mid-course of the Teles Pires River, Tapajós River basin, Brazil. Neotropical Ichthyology. 15.

Dehling, D. Matthias, P.J., Schaefer, H.M., Böhning-Gaese, K., and Schleuning, M, 2016. Morphology Predicts Species’ Functional Roles and Their Degree of Specialization in Plant– Frugivore Interactions. Proceedings of the Royal Society B: Biological Sciences 283, no. 1823:20152444.

Dumont, E.R., 1997. Cranial shape in fruit, nectar, and exudate feeders: implications for interpreting the fossil record. American Journal of Physical Anthropology: The Official Publication of the American Association of Physical Anthropologists, 102(2), pp.187–202.

Dumont, E.R., 1999. The effect of food hardness on feeding behaviour in frugivorous bats (Phyllostomidae): an experimental study. Journal of Zoology, 248(2), pp.219–229.

Fedorov A., Beichel R., Kalpathy-Cramer J., Finet J., Fillion-Robin J-C., Pujol S., Bauer C., Jennings D., Fennessy F.M., Sonka M., Buatti J., Aylward S.R., Miller J.V., Pieper S., Kikinis R. 3D Slicer as an Image Computing Platform for the Quantitative Imaging Network. Magnetic Resonance Imaging. 2012 Nov;30(9):1323-41. PMID: 22770690. PMCID: PMC3466397.

Fleming, T.H. and Kress, W.J., 2011. A brief history of fruits and frugivores. Acta Oecologica, 37(6), pp.521–530.

Fleming, T.H. and Kress, W.J., 2019. The ornaments of life: coevolution and conservation in the tropics. University of Chicago Press.

Freeman, P.W., 1988. Frugivorous and animalivorous bats (Microchiroptera): dental and cranial adaptations. Biological Journal of the Linnean Society, 33(3), pp.249–272.

Freeman, P.W., 1992. Canine teeth of bats (Microchiroptera): size, shape and role in crack propagation. Biological Journal of the Linnean Society, 45(2), pp.97–115.

González, N. & Vispo, C. 2002. Aspects of the diet and feeding ecologies of fish from nine floodplain lakes of the lower Caura, Venezuelan Guayana. Scientia Guaianae. 12:329–3.

Goulding, M. 1980. The fishes and the forest: explorations in Amazonian natural history. University of California Press, Berkeley.

Grubich, J.R., Huskey, S., Crofts, S., Ortí, G. and Porto, J., 2012. Mega-Bites: Extreme jaw forces of living and extinct piranhas (Serrasalmidae). Scientific Reports. 2, p.1009.

Horn, M.H., Correa, S.B., Parolin, P., Pollux, B.J.A., Anderson, J.T., Lucas, C., Widmann, P., Tjiu, A., Galetti, M. and Goulding, M., 2011. Seed dispersal by fishes in tropical and temperate fresh waters: the growing evidence. Acta Oecologica. 37(6), pp.561–577.

Howe, H.F., & Smallwood, J. Ecology of seed dispersal. Annual Review of Ecology and Systematics, 13, pp.201–228.

Huby, A., Lowie, A., Herrel, A., Vigouroux, R., Frédérich, B., Raick, X., Kurchevski, G., Godinho, A.L. and Parmentier, E., 2019. Functional diversity in biters: the evolutionary morphology of the oral jaw system in pacus, piranhas and relatives (Teleostei: Serrasalmidae). Biological Journal of the Linnean Society, 127(4), pp.722–741.

Huie, J.M., Summers, A.P. and Kolmann, M.A., 2019. Body shape separates guilds of rheophilic herbivores (Myleinae: Serrasalmidae) better than feeding morphology. Proceedings of the Academy of Natural Sciences of Philadelphia. 166(1), pp.1–15.

Jordano, P., Forget, P.M., Lambert, J.E., Böhning-Gaese, K., Traveset, A. and Wright, S.J., 2011. Frugivores and seed dispersal: mechanisms and consequences for biodiversity of a key ecological interaction.

Kinzey, W.G. and Norconk, M.A., 1990. Hardness as a basis of fruit choice in two sympatric primates. American Journal of Physical Anthropology, 81(1), pp.5–15.

Kinzey, W.G. and Norconk, M.A., 1993. Physical and chemical properties of fruit and seeds eaten by Pithecia and Chiropotes in Surinam and Venezuela. International Journal of Primatology, 14, pp.207–227.

Kolmann, M.A., Cohen, K.E., Bemis, K.E., Summers, A.P., Irish, F.J. and Hernandez, L.P., 2019. Tooth and consequences: Heterodonty and dental replacement in piranhas and pacus (Serrasalmidae). Evolution & Development. P.e12306.

Kolmann, M.A., Hughes, L.C., Hernandez, L.P., Arcila, D., Betancur-R, R., Sabaj, M.H., López-Fernández, H. and Ortí, G., 2021. Phylogenomics of piranhas and pacus (Serrasalmidae) uncovers how dietary convergence and parallelism obfuscate traditional morphological taxonomy. Systematic Biology, 70(3), pp.576–592.

Kolmann, M.A., Poulin, E., Rosen, J., Hemraj-Naraine, D. and Burns, M.D., 2024. Phenotypic convergence is stronger and more frequent in herbivorous freshwater fishes. Integrative and Comparative Biology, p.icae037.

Lambert, J.E., 1999. Seed handling in chimpanzees (Pan troglodytes) and redtail monkeys (Cercopithecus ascanius): Implications for understanding hominoid and cercopithecine fruit-processing strategies and seed dispersal. American Journal of Physical Anthropology: 109(3), pp.365–386.

Lucas, P.W. & Luke, D.A. 1984. Chewing It over: Basic Principles of Food Breakdown. In: Food Acquisition and Processing in Primates. Eds. David J. Chivers, Bernard A. Wood, Alan Bilsborough. Springer New York, NY pp. 283-301.. doi.org/10.1007/978-1-4757-5244-1

Nico, L.G. and Taphorn, D.C., 1988. Food habits of piranhas in the low llanos of Venezuela. Biotropica. 20(4), pp.311–321.

Nico, 1990. Trophic ecology of piranhas (Characidae: Serrasalminae) from savanna and forest regions in the Orinoco River basin of Venezuela. Unpubl. Ph.D. diss., Univ. of Florida, Gainesville.

Nico, L.G. & de Morales, M., 1994. Nutrient content of piranha (Characidae, Serrasalminae) prey items. Copeia. (2): 524–528.

Norconk, M.A. and Veres, M., 2011. Physical properties of fruit and seeds ingested by primate seed predators with emphasis on sakis and bearded sakis. The Anatomical Record: Advances in Integrative Anatomy and Evolutionary Biology, 294(12), pp.2092–2111.

Prudente, B.D.S., Carneiro-Marinho, P., Valente, R.D.M. and Montag, L.F.D.A., 2016. Feeding ecology of Serrasalmus gouldingi (Characiformes: Serrasalmidae) in the lower Anapu River region, eastern Amazon, Brazil. Acta Amazonica, 46(3), pp.259–270.

Rojas, D., Vale, A., Ferrero, V. and Navarro, L., 2012. The role of frugivory in the diversification of bats in the Neotropics. Journal of Biogeography, 39(11), pp.1948–1960.

Röpke, C.P., Ferreira, E. and Zuanon, J., 2014. Seasonal changes in the use of feeding resources by fish in stands of aquatic macrophytes in an Amazonian floodplain, Brazil. Environmental biology of fishes. 97(4): 401–414.

Sivault, E., McConkey, K.R., Bretagnolle, F., Sengupta, A., Lambert, J., Heymann, E., Herrel, A., and Forget, P., 2023. Can Body Mass and Skull Morphology Predict Seed and Fruit Ingestion Potential for Mammal Species? A Test Using Extant Species and Its Application to Extinct Species. Functional Ecology, 37, pp. 1504–1515.

Stiles, E.W. and White, D.W., 1986. Seed deposition patterns: influence of season, nutrients, and vegetation structure. Frugivores and seed dispersal, pp.45–54.

Thompson, A.W., Betancur-R, R., López-Fernández, H. and Ortí, G. 2014. A time-calibrated, multi-locus phylogeny of piranhas and pacus (Characiformes: Serrasalmidae) and a comparison of species tree methods. Molecular Phylogenetics and Evolution, 81, pp.242–257.

Valenta, K., Daegling, D.J., Nevo, O., Ledogar, J., Sarkar, D., Kalbitzer, U., Bortolamiol, S., Omeja, P., Chapman, C.A., Ayasse, M. and Kay, R., 2020. Fruit selectivity in anthropoid primates: size matters. International Journal of Primatology, 41, pp.525–537.

Vogel, E.R., Zulfa, A., Hardus, M., Wich, S.A., Dominy, N.J. and Taylor, A.B., 2014. Food mechanical properties, feeding ecology, and the mandibular morphology of wild orangutans. Journal of Human Evolution, 75, pp.110–124.

Wotton, D.M., Drake, D.R., Powlesland, R.G. and Ladley, J.J., 2016. The role of lizards as seed dispersers in New Zealand. Journal of the Royal Society of New Zealand, 46(1), pp.40–65.

